# Predicting functional networks from region connectivity profiles in task-based versus resting-state fMRI data

**DOI:** 10.1101/259077

**Authors:** Javier Rasero, Hannelore Aerts, Marlis Ontivero Ortega, Jesus M. Cortes, Sebastiano Stramaglia, Daniele Marinazzo

**Affiliations:** Biocruces Health Research Institute. Hospital Universitario de Cruces. E-48903, Barakaldo, Spain.; Faculty of Psychology and Educational Sciences, Department of Data Analysis, Ghent University, Henri Dunantlaan 2, B-9000 Ghent, Belgium; Neuroinformatics Department, Cuban Center for Neuroscience (CNeuro), La Habana, Cuba; Ikerbasque, The Basque Foundation for Science, E-48011, Bilbao, Spain.; Dipartimento di Fisica, Universitá degli Studi “Aldo Moro” Bari, Italy; Istituto Nazionale di Fisica Nucleare, Sezione di Bari, Italy May 12, 2018

## Abstract

Intrinsic Connectivity Networks, patterns of correlated activity emerging from “resting-state” Blood Oxygenation Level Dependent time series, are increasingly being associated to cognitive, clinical, and behavioral aspects, and compared with the pattern of activity elicited by specific tasks. We study the reconfiguration of the brain networks between task and resting-state conditions by a machine learning approach, to highlight the Intrinsic Connectivity Networks (ICNs) which are more affected by the change of network configurations in task vs. rest. We use a large cohort of publicly available data in both resting and task-based fMRI paradigms; by trying a battery of different supervised classifiers relying only on task-based measurements, we show that the highest accuracy is reached with a simple neural network of one hidden layer. In addition, when testing the fitted model on resting state measurements, such architecture yields a performance close to 90% for areas connected to the task performed, which mainly involve the visual and sensorimotor cortex, whilst a relevant decrease of the performance is observed in the other ICNs. On one hand, our results confirm the correspondence of ICNs in both paradigms (task and resting) thus opening a window for future clinical applications to subjects whose participation in a required task cannot be guaranteed. On the other hand it is shown that brain areas not involved in the task display different connectivity patterns in the two paradigms.

**Abbreviations**
ICN: Intrinsic Connectivity Network
BOLD: Blood Oxygenation Level
fMRI: Functional magnetic resonance imaging
VIS: Visual Network
SM: Somatosensory Network
VA: Ventral Attention Network
DA: Dorsal Attention Network
L: Lymbic Network
FP: Fronto-pariental Network
DMN: Default Mode Network
CER: Cerebellar Network
SUB: Subcortical Network
FSL: FMRIB Software Library
FLIRT: FMRIB's Linear Image Registration Tool
FNIRT: FMRIB's Non-Linear Image Registration Tool
HCP: Human Connectome Project
RF: Random Forest
SVM: Support Vector Machines
NN: Neural Network
ROC: Receiver Operating Characteristic
PR: Precision-Recall
TPR: true positive rate
FPR: false positive rate

## 1 Introduction

Functional magnetic resonance imaging (fMRI) has become a powerful tool to study brain dynamics with relatively fine spatial resolution. One popular paradigm, given the indirect and contrast-based nature of the method, is the block design, alternating task with passive rest. Task-associated brain activity can then be inferred by contrasting the level of BOLD signal between task and resting blocks in each voxel.

In a seminal study by Biswal *et al*. [Biswal et al., 1995], it has been shown that spontaneous low frequency fluctuations (< 0.1 Hz) in BOLD signal, present even when subjects are not performing any specific task, give rise to correlated patterns. In particular, after identification of seed regions in the sensorimotor cortex by a bilateral finger tapping task fMRI protocol, the authors found synchronous fluctuations of BOLD time courses between these seed regions and homologous areas in the opposite hemisphere. This finding demonstrated the existence of a sensorimotor network even at resting state. In addition to the sensorimotor network, several other functional networks that are activated in task designed experiments have since been identified during resting state, such as the dorsal and ventral attention networks, and the fronto-parietal control network [Fox et al., 2006, Vincent et al., 2008]. Among these functional networks, the most characteristic and ubiquitous is the default mode network (DMN) [Raichle et al., 2001], including the posterior cingulate cortex, precuneus and medial prefrontal cortex. Regions belonging to this network exhibit a deactivation when cognitive tasks are performed and an increase in activity during rest [Gusnard and Raichle, 2001].

In order to identify these Intrinsic Connectivity Networks (ICNs) emerging from resting-state measurements, the most common methods are seed based analysis and independent component analysis. In seed-based analysis, voxels within a region of interest (“seed region”) are selected and their average BOLD time course is correlated with that of all other voxels in the brain. Voxels showing a correlation with the seed region above a certain threshold are then considered to be functionally related to the region of interest. The main disadvantage of this approach, however, is that it requires *a priori* selection of seed regions. Independent component analysis (ICA), in contrast, is a model-free approach, requiring very few assumptions [Beckmann et al., 2005]. This method separates the BOLD time courses of all voxels into different spatial components and ensures maximum statistical independence among them [Damoiseaux et al., 2006]. Since this is a pure data-driven approach, in contrast to the “seed region” scenario, there is no clear link between the components found and the specific brain functions. As a result, component labeling might not be straightforward, especially at the individual subject level. Furthermore, the number of components to be retained is an arbitrary parameter to be provided, that might be fitted by a supplied criterion, such as the minimisation of a cost-sensitive function that allows an optimal match of the similarity reached by ICA predictions with respect to the observed dataset.

ICNs in resting state experimental designs can thus be regarded as regions with similar BOLD signal profiles. Therefore, different clustering methods can also be applied to explore the structure of whole-brain BOLD time series, in an attempt to identify functional networks on the basis of resting-state fMRI data (see [Cordes et al., 2002, Lee et al., 2012, Bellec et al., 2010, van den Heuvel et al., 2008] for the use of different unsupervised clustering methods). In order to solve the component labeling problem, supervised learning approaches can also be utilized, where the identification and prediction of these networks is accomplished after fitting the decision boundaries that separate each class by directly supplying the instance labels. In previous studies, a multilayer perceptron was found to be the optimal method to map the topography of ICNs in healthy young control subjects, after which it was applied to a small sample of patients undergoing surgical treatment for intractable epilepsy or tumor resection [Hacker et al., 2013, Mitchell et al., 2013]. The ability to reliably estimate functional networks from resting-state fMRI data would have important clinical implications, for example in neurosurgeons’ pre-surgical planning when patients are unable to cooperate with the task-based paradigm [Lee et al., 2016]. Furthermore, predictions from models uniquely trained on resting state measurements from healthy people have been demonstrated to robustly match activation profiles of pre-surgical populations that usually suffer from a great deal of variability [Jones et al., 2017].

The aim of the present study is to extend previous works looking at reconfiguration of brain networks between task and resting-state conditions, and to highlight the brain regions which are more affected by the change of network configurations in task vs. rest. We will quantify to which extent one can predict ICNs training in task-based paradigm and predict the ICN patterns emerging purely from resting-state fMRI data, using the optimal model as identified with task-based fMRI data. The development of a general framework for ICN identification using supervised machine learning methods could thus shed further light on the overlap between the patterns emerging from both task and resting data.

## 2 Materials and Methods

### 2.1 Subjects

We considered both resting-state and task-based fMRI data from 282 unrelated healthy subjects provided by the s900 release Human Connectome Project (HCP) [Essen et al., 2013]. All images were reconstructed using algorithm r227. This reconstruction algorithm directly performs the separation of the multi-band multi-slice in k-space, in contrast to a previous reconstruction algorithm (r177), where the separation occurred after transforming the fully acquired sampled data to frequency space along the read-out direction.

### 2.2 Data acquisition and preprocessing

Data were acquired on a customized Siemens 3T “Connectome Skyra” scanner, housed at Washington University in St. Louis, using a standard 32-channel Siemens receive head coil and a “body” transmission coil designed by Siemens specifically (Echo Planar Imaging sequence, Gradient-echo EPI, 1200 volumes, TR = 720 ms, echo time = 33.1 ms, flip angle = 52°, voxel size = 2 × 2 × 2mm^3^, field of view = 208 × 180mm^2^, 72 transversal slices).

Resting-state and task-based fMRI data were collected in two sessions. Each session consisted of two resting-state acquisitions of approximately 15 minutes each, where subjects were instructed to keep their eyes open, followed by task-based fMRI acquisitions of varying durations. For this study, we used the resting-state fMRI data acquired with Left-Right orientation from the first session, and task-based fMRI data with the MOTOR paradigm. In this task, participants were presented with visual cues that ask them to either tap their left or right finger, squeeze their left or right toe, or move their tongue. Each task volume image was obtained from 10 blocks of 12 seconds each, corresponding to 2 tongue movements, 4 hand movements (2 right and 2 left), 4 foot movements (2 right and 2 left), and 3 fixation blocks of 15 seconds. Each block was also preceded by a 3-second cue.

For this study, we used the preprocessed resting-state fMRI data as provided by HCP. In particular, the ICA-FIX pipeline [Glasser et al., 2013] was applied, which cleans the BOLD signal by removing noise components given by the ICA algorithm and that has been proved to increase the quality of the original data [Salimi-Khorshidi et al., 2014]. In the case of task data, such a denoised version was not yet available. Therefore, we downloaded the minimally preprocessed data and ICA-FIX processed it through the *hcp_fix* script that can be found in the HCP repository. This script performs a rigid-body head motion correction, high-pass temporal filtering, ICA decomposition of the data, FIX identification of the ICA components corresponding to artifacts and elimination of these components from the original data. Furthermore, we used FSL’s FNIRT command *applywarp* to transform both paradigm data from MNI space back to subjects’ native space through the appropriate matrix transformation that can be found for each subject in the HCP repository and FSL’s FLIRT to resample the native structural image at 2mm.

Next, we registered our volume data into the Shen parcellation atlas in the subjects’ native space, partitioning each subject’s brain in 268 functionally homogeneous and spatially coherent brain regions [Shen et al., 2013]. Functional connectivity matrices for each subject were then obtained by computing the Pearson correlation between the mean BOLD time course among all pairs of the 268 brain regions. A row (column) *i* of a given matrix thus represents the correlation map of the *i^th^* ROI with all other brain regions. Hence, these correlation maps describe the interactions of a brain node, which define the intrinsic connectivity networks (ICNs) assignment. Specifically, we considered seven cortical ICNs as proposed by Yeo and colleagues [Yeo et al., 2011] (visual (VIS), sensorimotor (SM), dorsal attention (DA), ventral attention (VA), limbic (L), fronto-parietal (FP), and default mode network (DMN)), and two more sub-networks for the sake of completeness, comprising subcortical (SUB) and cerebellar regions (CER). The number of regions per network in this atlas is 30, 25, 18, 22, 22, 23, 42, 48 and 38 respectively. ICN assignment to each of the 268 brain regions was performed by overlapping both Shen and Yeo atlas with a minimal 80% threshold. As a consequence, if the number of voxels within a Shen parcel exceeding this threshold belongs to one of Yeo’s ICNs, the parcel is assigned to that particular ICN.

Finally, for both task and resting data separately, the 282 resulting individual 268 × 268 Pearson correlation matrices were concatenated together row-wise to obtain *χ*_task_ and *χ*_rest_ super-matrices of 282 × 268 instances and 268 features each. Both these matrices are then later used in the subsequent machine learning analysis. Moreover, since self correlations, *i.e*. values of 1 in the Pearson connectivity matrices, would straightforwardly identify the ICN in a one-hot encoding fashion, we set these features to zero so that we guarantee that regions interactions to the whole brain are indeed the responsible for determining the ICN label.

### 2.3 Classification

Each correlation map was used to identity ICNs by means of four well known classification algorithms in machine learning: Quadratic Discriminant Analysis (QDA), Support Vector Machine (SVM) and Random Forest (RF) run under scikit-learn 0.18, an open source Python library that implements a wide range of machine learning tools which include preprocessing, cross-validation and classification algorithms [Pedregosa et al., 2011]; and a multilayer Neural Network (NN) using Keras in a Tensorflow backend, an API for deep learning written in Python that allows for a robust and easy implementation of neural networks with any kind of complexity desired [Chollet et al., 2015].

QDA is a linear classifier that stands out for being computationally friendly, inherently multiclass and very simple, with no hyperparameters to tune. In this classifier, the posterior label conditional probability functions *p*(*x*|*y*) are obtained through a likelihood taken as multivariate Gaussian distribution but, unlike in Linear Discriminant Analysis, covariances of the classes are not assumed to be equal, which lead to quadratic decision boundaries.

SVM is a maximum-margin classifier, aiming to find a hyperplane that maximizes the distance to the nearest points on each side representing each class (the so-called support vectors). It is specially suited for high dimensional problems, since it can linearly separate any dataset by going to a high dimensional space with the aid of a kernel function. For our analysis we used a linear kernel.

RF, on the other hand, is an ensemble of decision trees, which split the feature space according to the maximization of the Gini criterion, where each decision tree is trained on a random subset of examples and each splitting considers only a random subset of features, so that uncorrelated trees are obtained. By doing so, and averaging over all trees, one can reduce the variance and therefore avoid overfitting. For this study, 500 trees were considered.

Multilayer neural networks are able to fit a nonlinear function by passing the examples through its architecture while minimizing a loss function. The basic setup is a first layer with units equal to the number of features, a (few) hidden layer(s) and a final output layer that encodes the label information. The training is performed by updating the links (weights) among layers by back propagation in order to reduce the output error. We have explored several network architectures varying both depth and number of units per layer. Each layer has a *ReLu* activation except for the last one defining the example class which is a *Softmax* function. We used a Stochastic gradient descent optimizer with a learning rate of 0.01, a decay of 10^−6^ and a Nesterov momentum of 0.9. Likewise, in order to prevent overfitting we adopted a early stopping criterion on a %10 of the training dataset and a dropout rate of 0.2, which randomly sets this fraction rate of input units to 0 at each update during training time.

While MLP and RF can inherently map correlation maps into a 9-dimensional space, such an implementation is not as straightforward in SVM which is more suitable for binary classification. In order to address this issue, a common approach is to use a one-versus-all strategy, where one builds as many classifiers as class labels and trains each one taking one label as positive versus the rest being negative cases. Finally, classification of a test example is made by reporting the highest confidence score predicted after applying all classifiers.

We also applied such one-versus-all strategy to RF so that one can isolate the features which contribute most to the splitting in each classifier. Furthermore, when adopting the mentioned strategy, it is also important to take into account that the dataset will be unbalanced when building each classifier, so we have solved this by making both SVM and RF cost-sensitive, so that training an example from the majority class costs more than one of the minority class, weighted by the proportion of examples in both classes.

The best model was selected by comparing their global performance in a 5 times repeated 10-fold cross-validation scheme. Consequently, we first split the data into 10 equal size subsamples or folds such that one is used for testing while the rest fits the model. This process is then iterated for each of the k folds in order for them to be used once as test data. Furthermore, setting k=10 appears to be an optimal choice in terms of the trade-off between variance and bias of the classifier [Kohavi, 1995]. Second, the explained procedure is repeated 5 times shuffling the whole data each time, yielding then 5 random partitions of the original sample. The results from each of 10-fold cross-validation repetition can be averaged to assess stability of the classifier or otherwise merged all together (50 folds in our case) and averaged to produce a single estimation. In particular, we will adopt the latter one, except for model comparison, in which performance from each repetition is reported to address the aforementioned stability of the classifier.

On the other hand, given the nature of our data, where we have 268 observations for each subject, one could argue that these are correlated and therefore training and test datasets would not be totally independent if observations from the same subject fall in both subsamples. Such an effect is known as double-dipping and could lead to inflated association scores [Kriegeskorte et al., 2009]. We thus performed the 10-fold data partitions **based on subjects** so as to guarantee that observations from the same subjects are exclusively either in the training dataset or the test dataset.

We first obtained the best model by training and testing several algorithms using exclusively task data and second, the same model with the entire task fMRI as **training set** was then used to make predictions relying on resting fMRI data as **test set**. As such, one could infer whether intrinsic connectivity networks from a model fitted exclusively on task data could be recovered even when the subject is not able to perform the task.

Results are reported in several ways: (1) global accuracy *(i.e*. proportion of correctly classified instances); (2) a confusion matrix, where each row represents the instances in a predicted class and each column the examples in an actual class; (3) and ROC and Precision-Recall curves which exhibit model behaviour affected by different thresholds on the decision function predicted for each class. ROC curves represent the true positive (TPR) versus false positive rate (FPR), where in binary classification TPR = TP/P = #(pred==pos | pos)/(# pos) and FPR = FP/N= #(pred==pos | neg)/ (# neg), where # indicates number of examples. *Precision* is defined as the ratio of positive predictions that are indeed positive and *Recall* as the proportion of positive instances that are classified so.

### 2.4 Workflow

The different steps taken for the obtainment and use of the matrix of features within the machine learning algorithms analysis is shown in figure 1, which can be summarised as follows:

1. Formation of the matrix of feature vectors for both task and resting fMRI data.
2. 5 times repeated 10-fold cross-validation on **task** data to select the best predictive model. The data partition is based on subjects so that observations from the same subjects do not fall in both training and test data.
3. Results generalisation following the same partition procedure as before but now with the test samples containing only **resting** data whereas the model is still trained in task.

**Figure 1:**
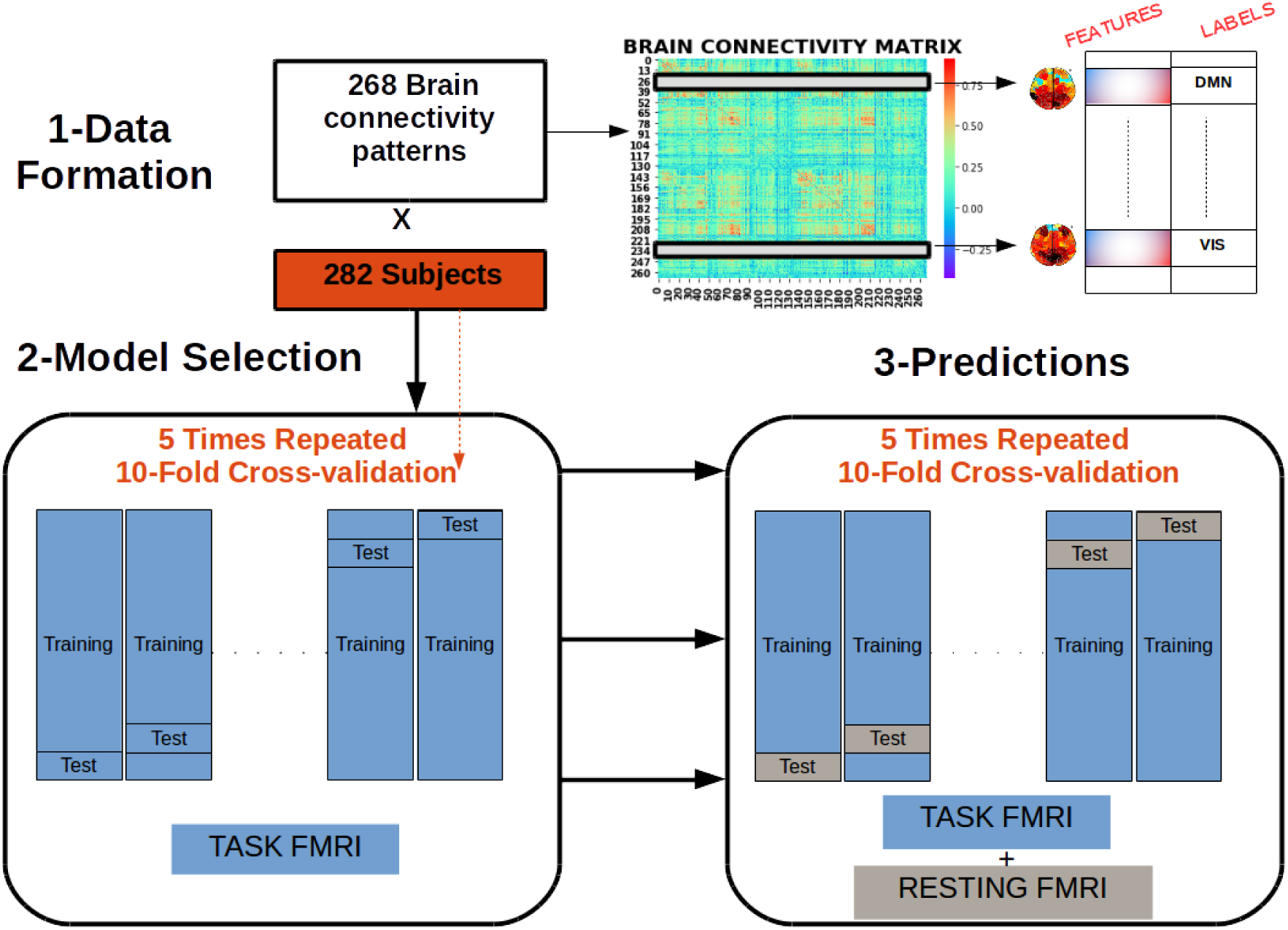
workflow describing how connectivity matrices for each subject were manipulated to obtain a matrix of features for both task and resting data separately and how both matrices were used subsequently to fit different machine learning algorithm and make predictions

## 3 Results

### 3.1 Performance on task fMRI

Global accuracy obtained after cross-validating the different methods described in the previous section can be seen in Figure 2.

**Figure 2:**
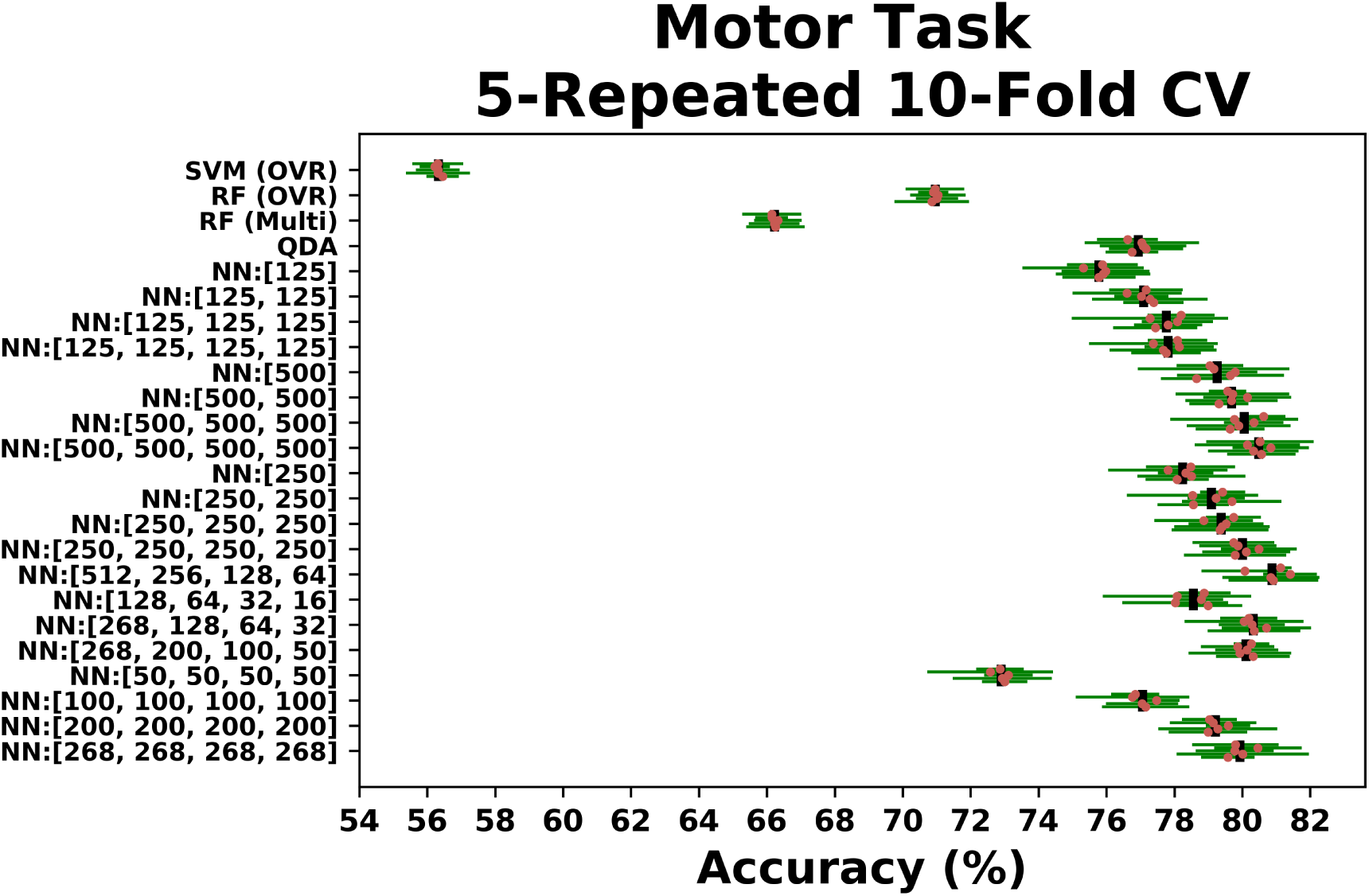
Averaged global accuracy within a 10-fold cross-validation scheme for 5 different shufflings of the data (orange points with green bars) for a set of neural network models (NN) with varying depth (length of the array in brackets in x-axis) and intermediate units (number within each component of the array in brackets in the x-axis), Random Forest in pure multi-class setting and Random Forest and Support Vector Machine both in a one-versus-rest scenario. Final averaged accuracy across the five shufflings is shown as a black vertical bar

First, we can see that the results across the different models are stable w.r.t different shufflings of the data. Second, neural networks outperform by far both Support Vector Machine and Random Forest algorithms, popular choices in literature when dealing with machine learning problems, and, as a consequence of the addition of complexity, they also improve in general the decent results of about 77% provided by Quadratic discriminant analysis classifier. Moreover, within the set of neural network models, there seems to be a tendency of better performance as the neural network becomes deeper, as well as the number of hidden units increases, which exhibits the demand for a higher number of parameters to define the decision boundaries separating each class. Notably, a decreasing complexity in architecture seems to capture better the intrinsic structure of the data, with a neural network with four hidden layers of 512, 256,128 and 64 units respectively, providing the best performance with a global accuracy of 81%. In the following, we will concentrate on the results given by this model.

Since we are dealing with a multi-class problem, it is important to calculate the classification performance of each ICN. This can been accomplished by means of the confusion matrix, that is depicted in Figure 3 for our best model. As one can see, all ICNs exhibit a good performance, specially for the visual, somatosensory-motor and dorsal attention where rates above 90% are achieved. The Limbic system however stands out of this behaviour, since its accuracy drops to approximately 66%, with a remarkable 18% of examples misclassified as subcortical regions.

**Figure 3:**
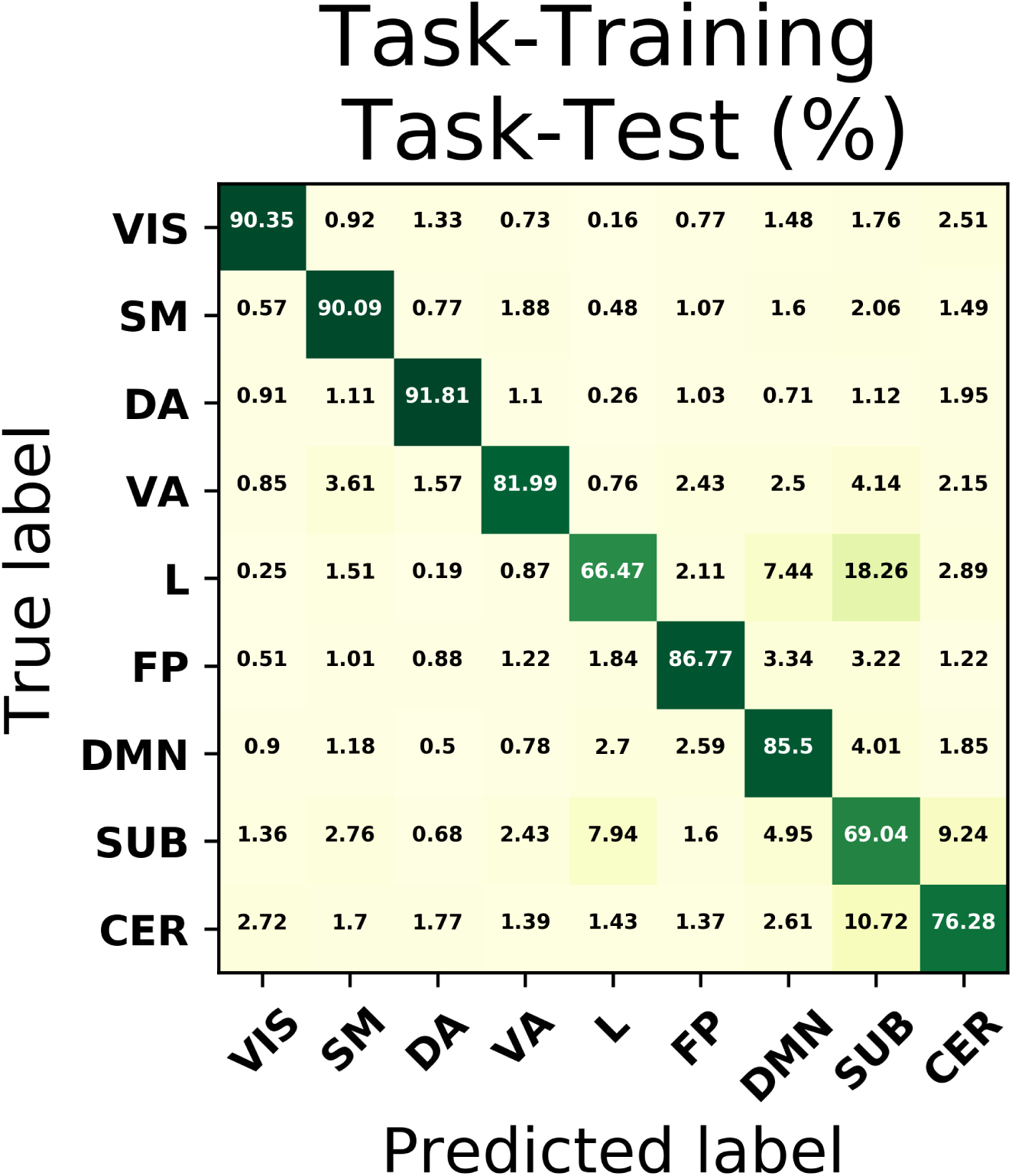
Confusion matrix for the best classification problem, which in this case corresponds to a NN with 4 hidden layer with 512, 256, 128 and 64 units

This difference in performance by each ICN might be addressed by looking at the similarity among them in more detail. In Figure 4, we show the Pearson cross-correlation matrix between pattern connectivities of the 268 nodes from the subjects-averaged connectivity matrix on the left, whereas on the right we depict the dispersion of the off-diagonal terms within each class (the bold boxes on the correlation matrix) in order to account for the intra-group similarity specifically. As it can be noticed, examples from the limbic and subcortical networks are more distant from each other, leading to an increase of similarity variance, which explains why the classifier struggles to build a decision function that distinguishes them efficiently. Likewise, the between-networks terms in the correlation matrix (those out of the black boxes) reflects the vicinity of subcortical regions to other network and it also shows that, yet having higher intra-group similarity, examples belonging to the Cerebellum are more misclassified than those of DMN ad FP, as a consequence of a smaller inter-group distance with other networks, especially with subcortical regions.

**Figure 4:**
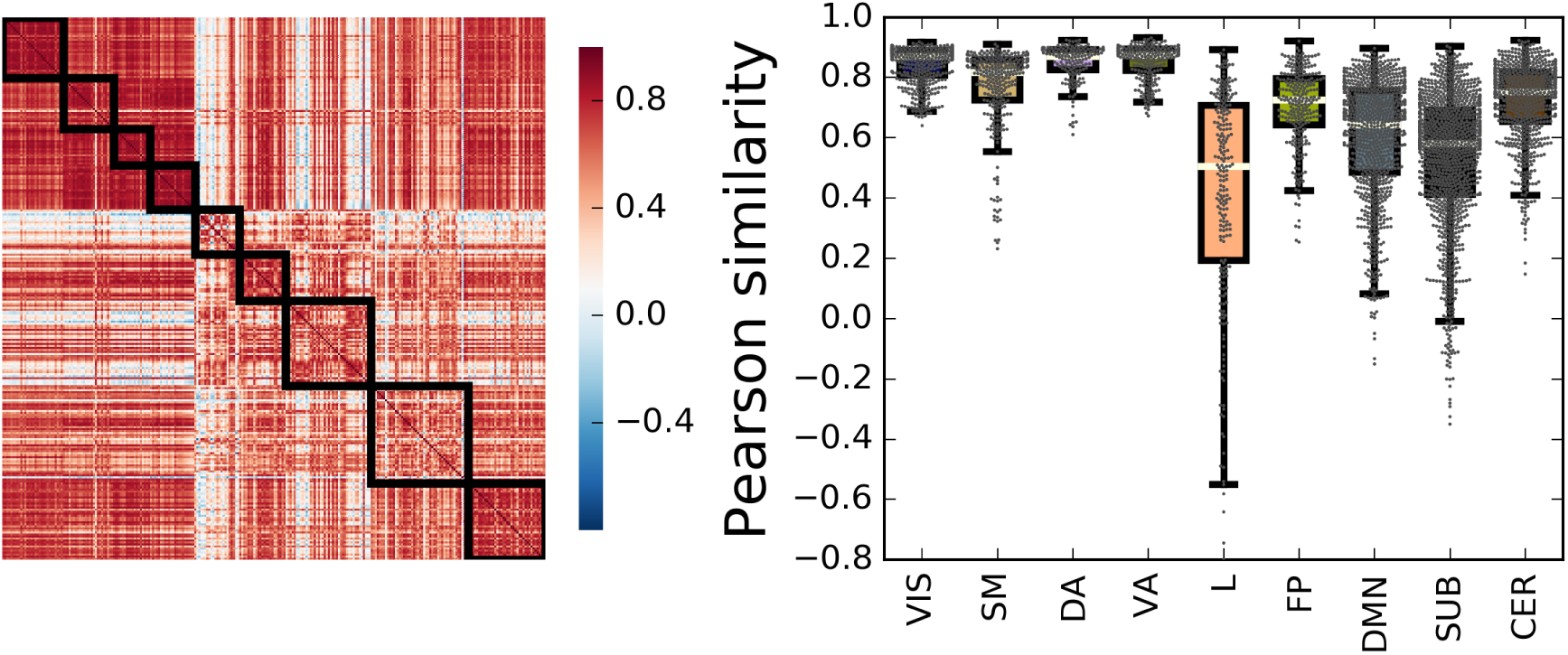
Representation of correlations amongst the 268 pattern connectivities from the subjects-averaged connectivity matrix. On the left, the full correlation matrix, ordered according to their group label, whose members’ interactions are outlined by a bold rectangle. On the right, the intra-group correlation distribution corresponding to the upper off-diagonal entries in each bold rectangle.

### 3.2 Generalisation to resting-state fMRI data

Next, we investigated how well the best neural network model generalised the results found so far when testing is performed on resting-state fMRI data from a model which was **only** trained using task-based data. In other words, we examined whether intrinsic connectivity networks can be successfully predicted in experiments where no active collaboration of the subjects is required.

Figure 5 shows the mean confusion matrix in a 5 times repeated 10-fold cross-validation based on subjects where the training has been performed after using the task-based fMRI dataset and testing on resting-state fMRI data. Results demonstrate that the same performance pattern exhibited in the previous section, whith a prominence from systems involved in the response to the stimuli triggered by the task protocol leading to high prediction rates in both VIS and SM network, with approximately 92% and 91% of sensitivity (also known as recall) respectively. Similarly, quick visual identification within the studied task to conditionally perform a specific movement encodes a bottom-up process which demands activation of ventral attention areas. Such areas are also well predicted by our fitted model with a 84% sensitivity. In contrast, dorsal attention areas attached to top-down stimuli drastically decrease their performance to 79%, with a moderate proportion being misclassified into the visual network, possibly reflecting some characteristic of the task paradigm in study. On the other hand, DMN regions whose activity negatively correlates with that of the other networks during rest exhibit also a decent performance of about 79%, which suggests that these areas maintain an intrinsic correlation that allows them to be optimally fitted by a model even during task. Finally, Limbic and Cerebellum systems involved in learning, memory and behaviour suffer from variability and hence are poorly replicated by our model.

**Figure 5:**
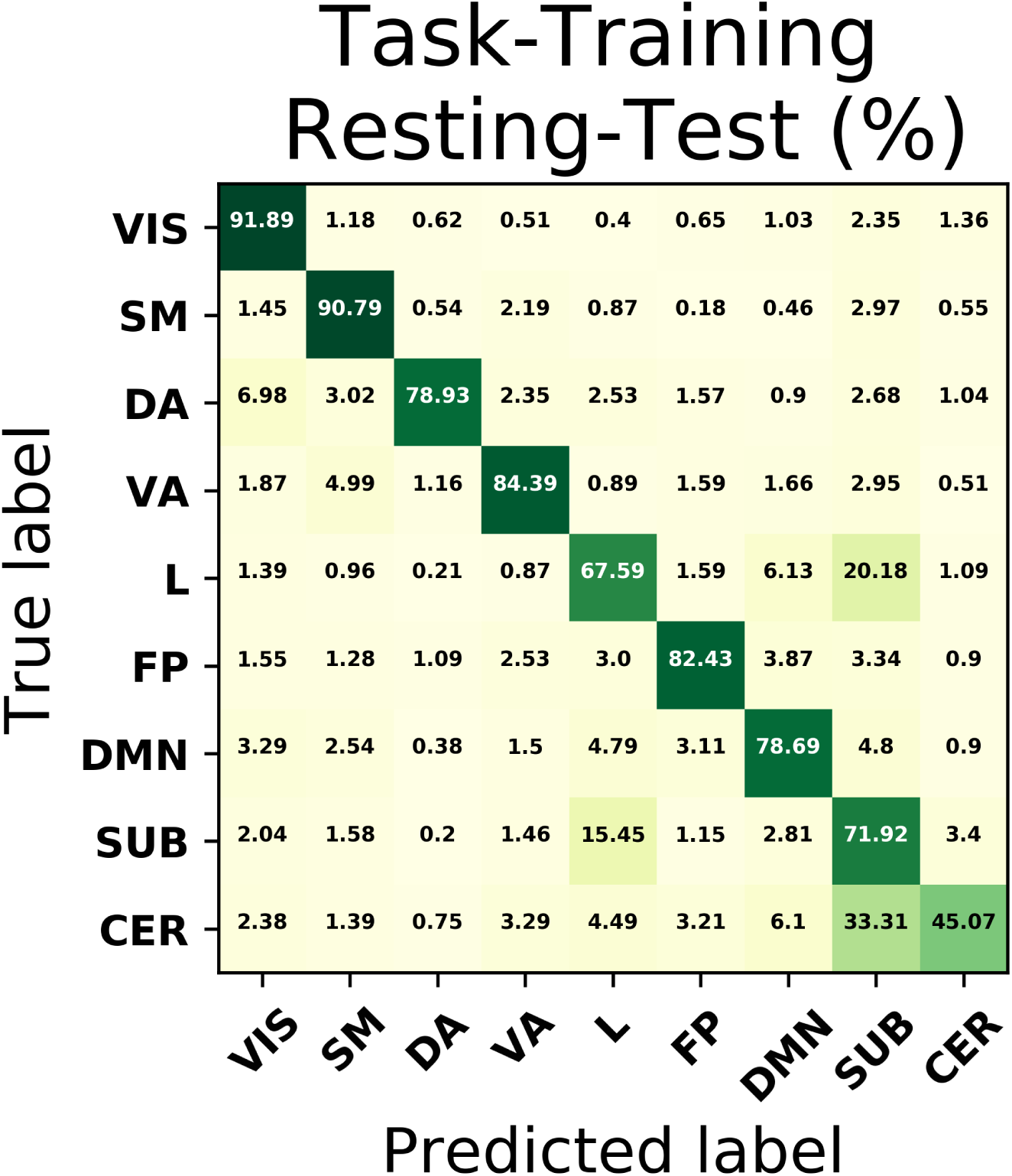
Confusion matrix for the best classification problem, which in this case corresponds to a neural network with four hidden layer of 512, 256, 128 and 64 units, using task-based fMRI data as training set and resting-state fMRI data as testing set.

The results obtained so far predict one *and only one* class for each instance, corresponding to the class assigned with the highest probability. Instead, we can directly look into this class probability prediction for each example and see how the model performs when changing the threshold of class assignment. We show this in Figure 6 and 7 through the ROC curves and the Precision-Recall curves. We can see that both representations describe the performance of the model when generalised to different settings, with most of the networks occupying large areas in both spaces and with a clear prominence from areas belonging to the SM and VIS networks. In addition, had we solely relied on results provided by the confusion matrix of figure 5, which represents a particular point of these curves, we would not have appreciated for example that the poorest performance of the Cerebellum network depended on a subpotimal threshold and could be improved beyond the performance of the Limbic and subcortical system.

**Figure 6:**
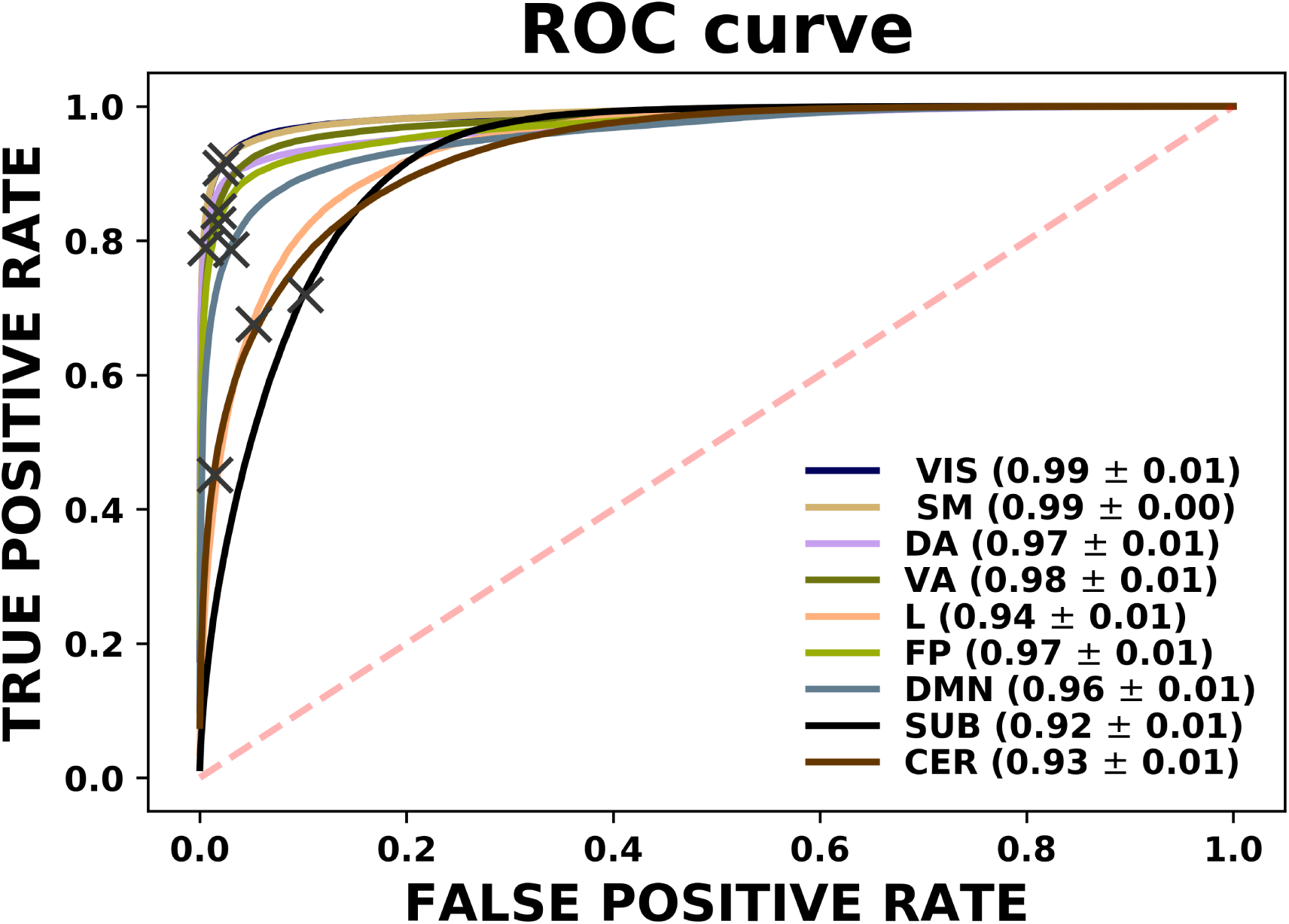
ROC curves for each class separately. The areas under these curves can be found in the legend located on the right side. Grey crosses display the model with the specific threshold yielding the results shown before. Those curves above the red dashed line represent exhibit a discriminating power

**Figure 7:**
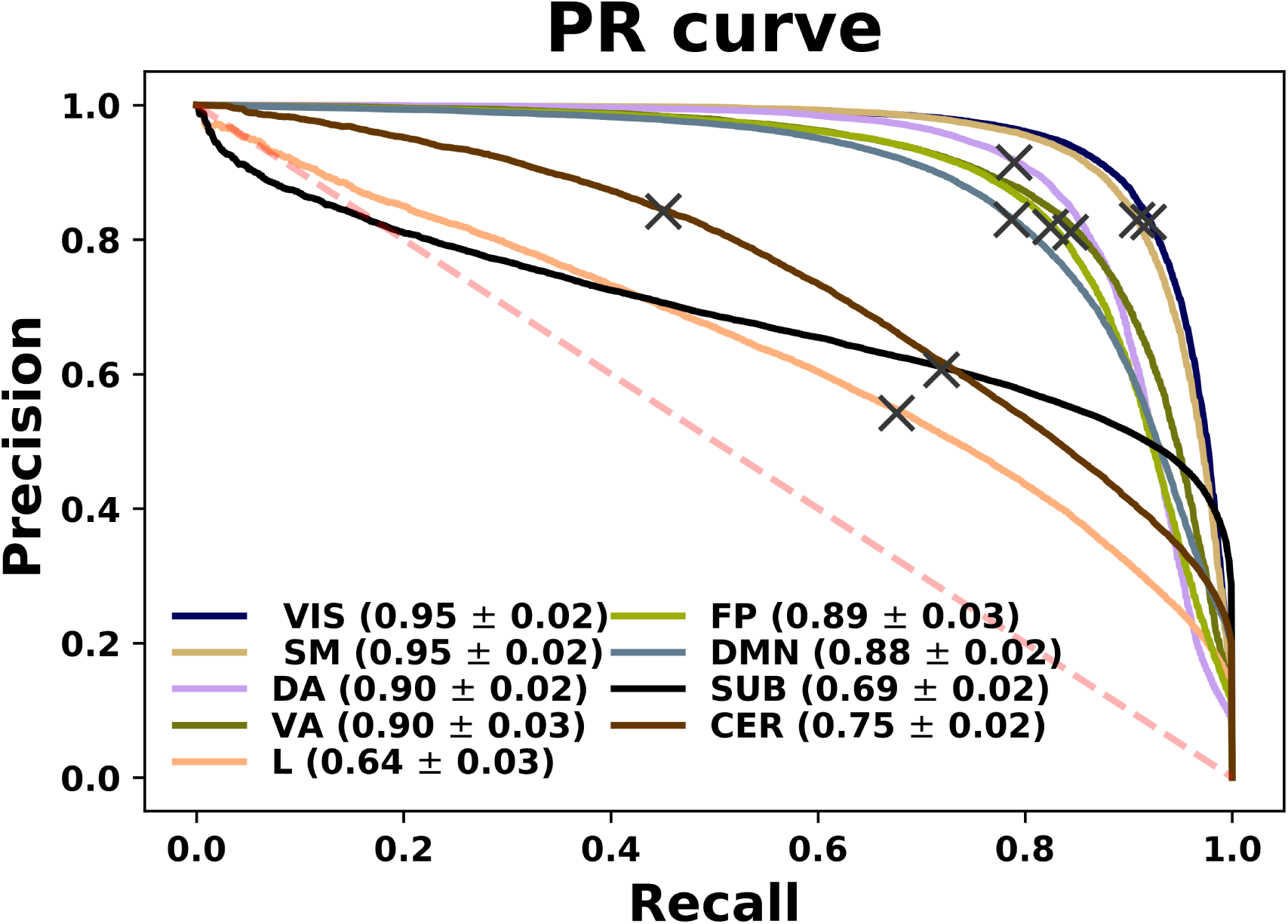
PR curves for each class separately. The areas under these curves can be found in the legend located on the right side. Grey crosses display the model with the specific threshold yielding the results shown before. Those curves above the red dashed line exhibit discriminating power.

Finally, it is reasonable to assume that performance from correlation patterns of different regions depend on the location of these latter in the brain. In order to address this issue, we show in Figure 8 the mean classification rate of each region across the different folds of the cross-validation procedure. This figure shows that, for instance, regions in the cerebellum display rather polarised results. Areas strictly in the right hemisphere within the posterior of the Crus Cerebellum I/II and lobes VI, VIIb and VIII have a successful accuracy greater than 80%, whereas anterior areas, lobes in the left hemisphere along with the vermis exhibit poorer performance than chance, being mainly mislabelled as subcortical regions. Cerebellum takes part in motor and attention tasks, where there is for example a clear asymmetry in the foot, hand, and tongue movement activation maps for intrinsic functional connectivity (left hemisphere) and task functional connectivity (right hemisphere) [Buckner, 2013]. On the other hand, being mainly involved in emotional and long-term cognitive functions, Limbic network is *a priori* the most detached system in relation with the task under exam here. Nonetheless, limbic regions located in the inferior and middle temporal gyrus and pole have decent classifying power, specially in the left hemisphere, whereas accuracy in orbital parts of the left frontal gyrus and rectus drastically drops below 50%. In addition, default mode network and fronto-parietal regions, which both yield a decent performance, exhibit interesting features. In general, they both have regions well above 90%, with higher incidence on the middle and posterior cingulum in both hemispheres and the left precentral and middle frontal gyrus, and poorly classifying parts depicted in yellow that correspond to the frontal gyrus orbital part, the olfactory cortex, rectus, hippocampus, calcarine and precuneus in the left hemisphere for default mode networks nodes; and the anterior and middle cingulum for both networks.

**Figure 8:**
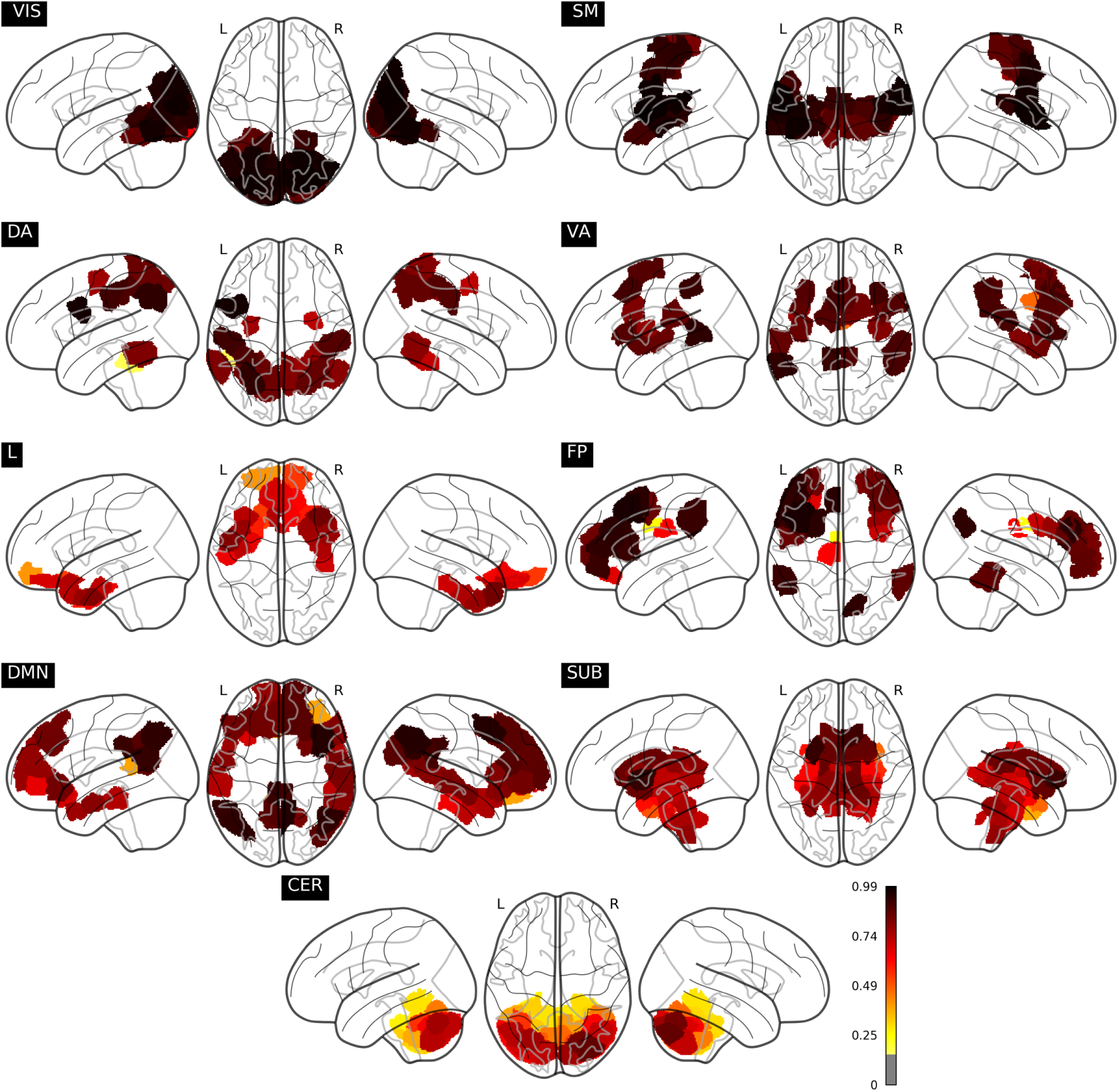
Mean classification accuracy of each node across all subjects contained in the test set in each fold of the cross-validation scheme.

## 4 Discussion

We have fitted different models on the whole-brain functional connectivity patterns obtained from the Pearson correlation matrices from a large population of healthy subjects, and quantify the success of the prediction of the organization in Intrinsic Connectivity Networks. We obtained the highest accuracy with a neural network of four hidden layers and 512, 256, 128 and 64 intermediate units each. Given the models trained, this result suggests that the complexity of the target function plays an important role with a demand of going deep in architecture, but also that mapping to decreasing numbers of intermediate units seems to better capture the inherent properties of the data, in accordance with the behavior of other well known machine learning algorithms such as autoencoders and convolutional neural networks. This behaviour could be even more evident if we considered a representation at the voxel level due to the increase of dimensionality, and therefore more complex models would likely be required. Moreover, well-established algorithms such as Random Forest (RF) and Support Vector Machines (SVM) were not capable of reaching a performance similar to the one obtained by neural networks. Two hypotheses might be formulated to explain this phenomenon. First, RF splits the space taking a subset of features, so this model fails to capture the optimal splitting out of the correlation maps since one would expect the whole pattern to be important. Second, both SVM and RF in general find it more challenging to build the boundary decision in high dimensional multi-class settings. On the other hand, the benefits of neural networks, that are well known for their capability to model accurately any arbitrary function [Lin et al., 2017], allow to improve the decent performance provided by the other attempted classifier QDA, suggesting that observations can be to some extension approximated by a multivariate Gaussian function.

Following the findings of a previous study [Smith et al., 2009], it is remarkable how accurately intrinsic connectivity networks can be identified using only task data, which reflects the high degree of correspondence between both modalities. Not only could visual and sensorimotor regions be clearly represented by both paradigms, as expected, but also other functional networks turned out to have correlation maps which lie within well defined decision boundaries. For example, time series of default mode network regions, though having lower participation during cognitive tasks, when cross-correlated with the average time series of all other brain regions, demonstrated that their integration and participation make them differentiable (in the end, Pearson correlation is unaltered by a change of scale). In contrast, the limbic system showed the worst performance, with limbic regions often miscategorised as belonging to the subcortical network. This was caused by some areas of both networks showing a similar pattern of activity, forcing the classifier to lean towards the subcortical network, the majority class over the limbic group.

The connection of task-evoked experiments with resting-state becomes even clearer if one looks at the results presented in figures 5, 6 and 7, where VIS and SM networks in particular (which are mainly involved in the MOTOR task), are recovered exclusively from resting data with great accuracy. This apparent correspondence between both modalities is being thoroughly investigated. In a seminal work, Smith *et al*. [Smith et al., 2009] demonstrated the existence of a high similarity between the ICA-analysis results extracted independently from a resting-state fMRI dataset consisting of 36 healthy subjects and activation maps from BrainMap database of functional imaging studies across nearly 30,000 human subjects. This suggests that the brain at rest is continuously active so as to be a composition of all the inherent possible tasks, such that a model trained based on a specific task will emerge naturally as one of its components when tested on resting state. On the other hand, in [Finn et al., 2015] an exhaustive study of this was carried out showing that the same networks that mostly discriminate individuals were also most predictive of cognitive behaviour. In [Tavor et al., 2016] prediction of activation maps by resting-state fMRI were overlapped with maps used to fit a wide range of task-based models and demonstrated that individual differences in brain response are inherently linked to the brain itself rather than to a specific manifestation given a certain task. Nonetheless, our work rather embraces the spirit of [Mitchell et al., 2013] in terms of the methodology used. We have aimed at successfully building the decision surfaces that separate the different ICNs by performing an exhaustive machine learning algorithm search in a large cohort of subjects. It is also worthwhile to stress that individual task-rest correspondence and between-subject reproducibility of patterns are different and possibly orthogonal problems, and that the cognitive relevance of these networks is mainly expressed on the subject-specific level [Kong et al., 2018].

Regarding the results obtained, one might be tempted to take the outcome of the ROC curves in Figure 6 as outstanding, where all the classes exhibit a ROC area greater than 0.9. However, as noted in [Davis and Goadrich, 2006], plotting ROC curves separately for each label might not be appropriate in multi-class settings where negative examples far exceed the number of positive instances. In this case *FP* ≪ *TN*, so changes in the number of false positives hardly affect the False Positive Rate, which continues to hold small and therefore the area obtained is usually large. As a consequence, it is better to use Precision-Recall curves, which concentrate only on the positive class and therefore do not suffer from these issues. As it can be seen in Figure 6, the areas found go more in concordance with the findings discussed previously, where the VIS and SM networks seem to be identified most accurately. In addition, we can see two interesting features from carefully inspecting these curves. Firstly, having both VIS and SM systems the largest areas, their observed thresholds (depicted in the figures as grey crosses) lie on a slightly non-optimal point on their respective curve. If we look at their performance in the diagonal of the confusion matrix of Figure 5, which is nothing less than the Recall measure, we therefore notice that this is accompanied by a loss of precision. Secondly, even though CER network seemed to behave extremely poor when testing the model on resting fMRI, we can see that this is an effect of the threshold used. In fact, when we vary this quantity, we can see that this group performs much better than the Limbic and Subcortical systems. Finally, it is also remarkable how well areas in DMN can be predicted from a model that was trained exclusively on task data, during which its activation would not be expected. One possible explanation might be that the functional connectivity within this network has a precise mapping with its structural connectivity [Greicius et al., 2009] and therefore, the integration in DMN might preserve across different tasks.

### Limitations

The interpretation of the results of the present study should be seen at the light of the following limitations. These are mainly related to the construction of the matrix of features and the definition of the labels to be classified.

First, the use of the correlations pattern to the whole brain of a given node as features that determine the ICN of that node should be treated carefully. In particular, setting all the terms but the self-correlation in the feature vector to zero, the resulting vector would directly provide the ICN label of the node with a 100 % of accuracy since each node vector would be unique and orthogonal to the rest and therefore classification would not be needed whatsoever. However, our hypothesis is that ICN assignment should be predicted by the whole pattern connectivity of the node. As a consequence, and given that we are limited by the fixed dimensions of the feature vectors (these can not change across subjects), we set self-correlations values to zero so as to diminish their effect and allow the rest terms to truly determine the label of each example.

Second, the labels used to fit the model are not fully and uniquely defined since they are threshold dependent when matching Shen with Yeo’s atlas. As a result, this spatial variability might somewhat blur the connectivity maps and therefore reduce the performance.

## 5 Conclusion

Successfully delineating intrinsic connectivity networks of the brain is of key importance to fully understand its behaviour and the patterns emerging during evoked activity (and vice-versa). Multivariate methods in machine learning turn out to be promising techniques in successfully identifying ICN networks from both task and resting-state fMRI data. As usual, the selection of the optimal algorithm is a crucial step. Obviously the optimal algorithm depends on the complexity of the problem. In our case, we have seen that a simple neural network with just one layer works the best. Whether a higher resolution or different parcellation might require to go deeper calls for future investigation.

As mentioned, the accurate prediction of the different baseline functions of the brain even when the model has been fitted using a different protocol (in our case a visual-tapping task) might be relevant for important future applications in clinical neuroscience. Localization of functions that are questionable due to the incapability of subjects to perform specific tasks could easily arise in this proposed scenario and could therefore help clinicians isolate the areas demanding a correct treatment for subject recovery. Moreover, future studies using the strategy followed in this work can address how brain regions in patients who have undergone a surgical operation change their connectivity pattern and therefore specialise in performing new tasks.

### Code

Code for downloading and preprocessing of the data used in this work, and replication of results and plots is available at https://github.com/jrasero/Predicting-icns

## Acknowledgment

JR acknowledges financial support from the Minister of Education, Language Policy and Culture (Basque Government) under Doctoral Research Staff Improvement Programme.

Data were provided [in part] by the Human Connectome Project, WU-Minn Consortium (Principal Investigators: David Van Essen and Kamil Ugurbil; 1U54MH091657) funded by the 16 NIH Institutes and Centers that support the NIH Blueprint for Neuroscience Research; and by the McDonnell Center for Systems Neuroscience at Washington University.

